# Host-specific subtelomere: Genomic architecture of pathogen emergence in asexual filamentous fungi

**DOI:** 10.1101/721753

**Authors:** Xiaoqiu Huang

## Abstract

Several asexual species of filamentous fungal pathogens contain supernumerary chromosomes carrying secondary metabolite (SM) or pathogenicity genes. Supernumerary chromosomes have been shown in *in vitro* experiments to transfer from pathogenic isolates to non-pathogenic ones and between isolates whose fusion can result in vegetative or heterokaryon incompatibility (HET). However, much is still unknown about the acquisition and maintenance of SM/pathogenicity gene clusters in the adaptation of these asexual pathogens to their hosts. We investigated several asexual fungal pathogens for genomic elements involved in maintaining telomeres for supernumerary and core chromosomes during vegetative reproduction. We found that in vegetative species or lineages with a nearly complete telomere-to-telomere genome assembly (e.g. *Fusarium equiseti* and five *formae speciales* of the *F. oxysporum* species complex), core and supernumerary chromosomes were flanked by highly similar subtelomeric sequences on the 3’ side and by their reverse complements on the 5’ side. This subtelomere sequence structure was preserved in isolates from the same species or from polyphyletic lineages in the same *forma specialis* (f.sp.) of the *F. oxysporum* species complex. Moreover, between some isolates within *F. oxysporum* f.sp. *lycopersici*, the mean rate of single nucleotide polymorphisms (SNPs) in a supernumerary chromosome was at least 300 times lower than those in core chromosomes. And a large number of HET domain genes were located in SM/pathogenicity gene clusters, with a potential role in maintaining these gene clusters during vegetative reproduction.

## Introduction

Asexual filamentous fungi lack genome-wide homologous recombination, a significant source of genetic variability in sexual filamentous fungi. Several asexual species of filamentous fungal pathogens contain supernumerary chromosomes (also known as accessory, conditionally-dispensable, or lineage-specific chromosomes), which are present in some but not all isolates (strains) of the species (Covert 1998). Supernumerary chromosomes carry secondary metabolite (SM) or pathogenicity gene clusters that are present in some isolates but absent from closely related isolates of the same or different species (Ma et al. 2010; Rep and Kistler 2010; Croll and McDonald 2012; Raffaele and Kamoun 2012). SM/pathogenicity gene clusters are also found in core chromosomes; they tend to be localized closer to the ends of core chromosomes (subtelomeres) (Ma et al. 2010; Wiemann et al. 2013; Dong et al. 2015; Niehaus et al. 2017). In several *Fusarium* species, supernumerary chromosomes are enriched for transposons and are highly variable among isolates (Ma et al. 2010; Huang et al. 2016; Vanheule et al. 2016; Vlaardingerbroek et al. 2016b). Another connection between core subtelomeres and supernumerary chromosomes is that they are enriched with a silencing histone modification, trimethylation of lysine 27 on histone H3 protein subunit (H3K27me3) (Galazka and Freitag 2014). Supernumerary chromosomes have been shown in *in vitro* experiments to transfer between vegetatively incompatible isolates or to transfer from a pathogenic isolate to a non-pathogenic isolate (Horizontal Chromosome Transfer or HCT) in asexual filamentous fungi (He et al. 1998; Akagi et al. 2009; Ma et al. 2010; Vlaardingerbroek et al. 2016a; van Dam et al. 2017). A phylogenetic analysis of core genes and supernumerary effector genes suggests horizontal transfer of these effector genes (van Dam et al. 2016). Thus, an important question is whether HCT occurs frequently in nature. Another one is whether there is homologous recombination between core and supernumerary chromosomes over their subtelomeres, which, combined with HCT of supernumerary chromosomes, allows SM/pathogenicity gene clusters to arise from core chromosomes of one lineage and to end up in core chromosomes of another one. This raises more questions in vegetative reproduction. One is whether the structure of core and supernumerary sub-telomeres can shed light on the expression of SM/pathogenicity gene clusters in subtelomeric heterochromatin. Another one is how these gene clusters are prevented from being lost from core chromosomes, as they are beneficial in certain environments, but not essential.

In the fission yeast *Schizosaccharomyces pombe*, adjacent to the end of the chromosome is the subtelomere containing species-specific homologous DNA sequences, whose functions include gene expression and chromosome homeostasis (Tashiro et al. 2017). In the budding yeast *Saccharomyces cerevisiae* and several filamentous fungi, the subtelomeres contain a helicase-like gene (Louis 1995; Sánchez-Alonso and Guzman 1998; Gao et al. 2002; Inglis et al. 2005; Rehmeyer et al. 2006). A DNA helicase regulates mutually exclusive expression of virulence genes in the subtelomeres of *Plasmodium falciparum* via heterochromatin alteration (Li et al. 2019). In the asexual pathogen *Fusarium oxysporum*, SM/pathogenicity gene clusters (facultative heterochromatin) are marked by histone H3 lysine 27 trimethylation (H3K27me3), while euchromatin is marked by histone H3 lysine 4 dimethylation (H3K4me2) (Fokkens et al. 2018). Because subtelomeres contain repetitive DNA, their DNA sequences are difficult to reconstruct by genome assembly programs, so they are sparsely represented in databases of whole genome sequence data (Wu et al. 2009). Only in *Colletotrichum higginsianum* isolate IMI 349063, was it reported that the supernumerary and core chromosomes possessed highly similar subtelomeric sequences containing a helicase-like gene (Dallery et al. 2017). This finding motivated us to investigate whether this subtelomeric structure is preserved within the same species or *forma specialis* so as to form its own unique type of subtelomeric sequences, and how those sequences arise.

Filamentous fungal subtelomeres contain repetitive DNA, which may be controlled by several genome defense mechanisms including Repeat-Induced Point mutation (RIP) (Cambareri et al. 1989; Gladyshev 2017). Because RIP operates during the sexual cycle in filamentous ascomycetes, fungi currently undergoing sexual reproduction are unlikely to possess intact subtelomere domains; for example, the presence of nucleotide changes indicative of RIP was shown on a family of highly similar telomere-associated helicases in the fungus *Nectria haematococca* Mating Population VI (MPVI) (Coleman et al. 2009), and no repetitive DNA or helicase was found near the ends of chromosomes in the sexual filamentous fungus *Neurospora crassa* (Wu et al. 2009). Thus, a relevant question is whether repetitive DNA in the subtelomeres of an asexual fungal pathogen (with a strong RIP system) was acquired during the asexual cycle or during the speciation of the asexual pathogen.

We attempted to understand the maintenance of SM/pathogenicity gene clusters in vegetative reproduction by exploring its connection to the genetic basis of vegetatively growing cells (in filamentous fungi) to distinguish self from nonself (allorecognition). During the vegetative growth phase, allorecognition can result in vegetative or heterokaryon incompatibility (HET) following fusion of genetically different cells, which disrupts growth and causes cell death (Glass and Dementhon 2006; Paoletti and Saupe 2009). However, heterokaryon incompatibility is suppressed following conidial anastomosis tube (CAT) fusion between vegetatively incompatible strains of *C. lindemuthianum* (Ishikawa et al. 2012), suggesting that HCT may result from CAT fusion (Manners and He 2011). Heterokaryon incompatibility involves a protein partner with a HET domain as a trigger of programmed cell death (Paoletti and Clavé 2007). A large number of HET domain genes were found in the genomes of asexual filamentous fungal pathogens, for example, 231 and 324 in two isolates of *Pyrenochaeta lycopersici* (Dal Molin et al. 2018). HET domain genes possess some characteristics of pathogenicity islands: they are highly variable between closely related isolates from different incompatibility groups, but they display trans-species polymorphism, where a HET domain gene from an isolate is more similar to one from an isolate of a different species than to ones from isolates of the same species (Muirhead et al. 2002). HET domain genes in three filamentous fungal species also exhibit a trend toward clustering near the ends of chromosomes (Zhao et al. 2015). The origins and functions of many HET domain genes in filamentous fungi remain unknown (Paoletti and Clavé 2007; Smith and Lafontaine 2013). A vexing question is whether HET domain genes tend to be found in SM/pathogenicity gene clusters and prevent them from being lost from core chromosomes.

Recently, long-read sequencing such as Single Molecule, Real-Time (SMRT) Sequencing has resulted in genome assemblies that span repetitive elements and complex regions of lengths up to 30 kb. In this paper, we address the above questions by analyzing SMRT-based genome sequences of asexual fungal pathogens. This analysis helps shed light on a number of molecular mechanisms that work together to acquire and maintain SM/pathogenicity gene clusters in these genomes.

## Results

### *F. oxysporum* f.sp. *lycopersici*

A genome assembly of *F. oxysporum* f.sp. *lycopersici* (Fol) race 3 isolate D11 consisted of 10 chromosomes and 29 contigs (Henry et al. 2019). We identified 16 subtelomeres in the chromosomes and 11 subtelomeres in the contigs, where the subtelomere of a chromosome or contig is a longest terminal region that ends in a telomeric repeat and is highly similar to terminal regions elsewhere. All 16 chromosome subtelomeres (in reverse orientation for 5’ ones and in forward orientation for 3’ ones) were at 99.89-99.98% identity over a length of 10,250-10,809 bp to one of them (the 3’ subtelomere of chromosome 1 in forward orientation), and so were 7 of the 11 contig subtelomeres. Among these were the 5’ subtelomere of supernumerary pathogenicity chromosome 14, and the 3’ subtelomere of supernumerary contig 38, which was highly similar to supernumerary chromosomes 3 and 6 in isolate Fol4287 (Ma et al. 2010). However, the remaining 4 contig subtelomeres, each with an extra section of 0.6 to 6.2 kb positioned around 7 kb from the telomeric end, were less similar to the above group of common subtelomeres. These 4 contig subtelomeres were divided into three different types. A first of them, the 3’ subtelomere of contig 41, was at 98.48% identity over 11,085 bp to a second of them, the 3’ subtelomere of contig 9, which was at 99.99% identity to a third of them, the 5’ subtelomere of contig 15 (in reverse orientation); so the second and third were of the same type. The fourth of them, the 5’ subtelomere of contig 16, was more similar to the third than the first; an alignment of the fourth and third contained a block of two different regions of 6,169 bp starting at 7,112 bp in contig 16 and of 712 bp starting at 7,111 bp in contig 15. Three of these 4 contigs, contigs 9, 15 and 41 of lengths 49, 514 and 25 kb, were supernumerary; the other one, contig 16 of 1,919 kb, was a core contig (see HET domain genes below). This core contig had a common 3’ subtelomere but a unique 5’ subtelomere containing an extra section of 6 kb.

To see whether these subtelomeres were present in other isolates, we selected 6 isolates from this *forma specialis*: Fol4287 (race 2), Fol007 (race 2), Fol014 (race 1), Fol026 (race 3), Fol069 (race 2), and Fol072 (race 2); and a non-pathogenic isolate: Fo47. The first 4 isolates were from the same lineage, and each of the other 3 isolates was from a different lineage (van Dam et al. 2016). Short reads from each of these isolates were mapped onto the Fol D11 genome assembly as a reference. Because the D11 subtelomeres were highly similar, no short reads could be uniquely mapped onto them. To fix this problem, short reads from each isolate were mapped onto the 3’ subtelomere of D11 chromosome 1 as a reference (see Methods). The estimated copy number of the D11 chromosome 1 subtelomere was 25 in Fol4287, 28 in Fol007, 32 in Fol014, 35 in Fol026, was 31 in Fol069, 34 in Fol072, and 0 in Fo47. The subtelomere was completely covered by short reads from each of the 6 pathogenic isolates, with at most 2 single nucleotide polymorphisms (SNPs) between isolate D11 and every other pathogenic isolate over the subtelomere. The mapping process was repeated for each of the 3 types of D11 contig subtelomeres, There was more variation between isolate D11 and the other pathogenic isolates in the 3 types of contig subtelomeres, which, for example, were not covered by any short reads from isolate Fol4287 at the border to the extra section of bases (see above).

Next we checked whether supernumerary D11 chromosome 14 and contig 38 were present and conserved in isolates Fol069 and Fol072, which were not in the same lineage as isolates D11 and Fol4287, as mentioned above. For each of isolates Fol4287, Fol069 and Fol072, short reads from the isolate were mapped onto the Fol D11 genome assembly as a reference, the mapped reference was partitioned into disjoint windows of maximum lengths such that each position in the window was covered at a depth greater than or equal to *adc/wca*, and each window of at least length *wlen* was selected to calculate its SNP rate, where *adc* is the average depth of coverage over the reference genome for the isolate, and *wlen* and *wca* are parameters. Note that windows were introduced to select regions of the reference that were unique and less variable between Fol D11 and the other isolates. The parameters *wlen* and *wca* were set to 10,000 bp and 5, respectively, to ensure that there were a sufficient number of windows for each isolate (Table 1). Supernumerary D11 chromosome 14 and contig 38 were present in each isolate, based on their percent coverage by short reads from the isolate (Table 1), which was the percentage of the chromosome or contig covered at a depth greater than or equal to *adc/wca*. And they were more conserved than the core chromosomes between Fol D11 and each of isolates Fol069 and Fol072, by the weighted mean of the SNP rates of windows in each of them in comparison to that in the genome (Table 1), where the weight for each rate was the size of its window divided by the sum of all window sizes. This was unexpected, as supernumerary chromosomes were much more variable than core chromosomes among isolates from the same species (Huang et al. 2016; Vlaardingerbroek et al. 2016b).

**Table 1.**
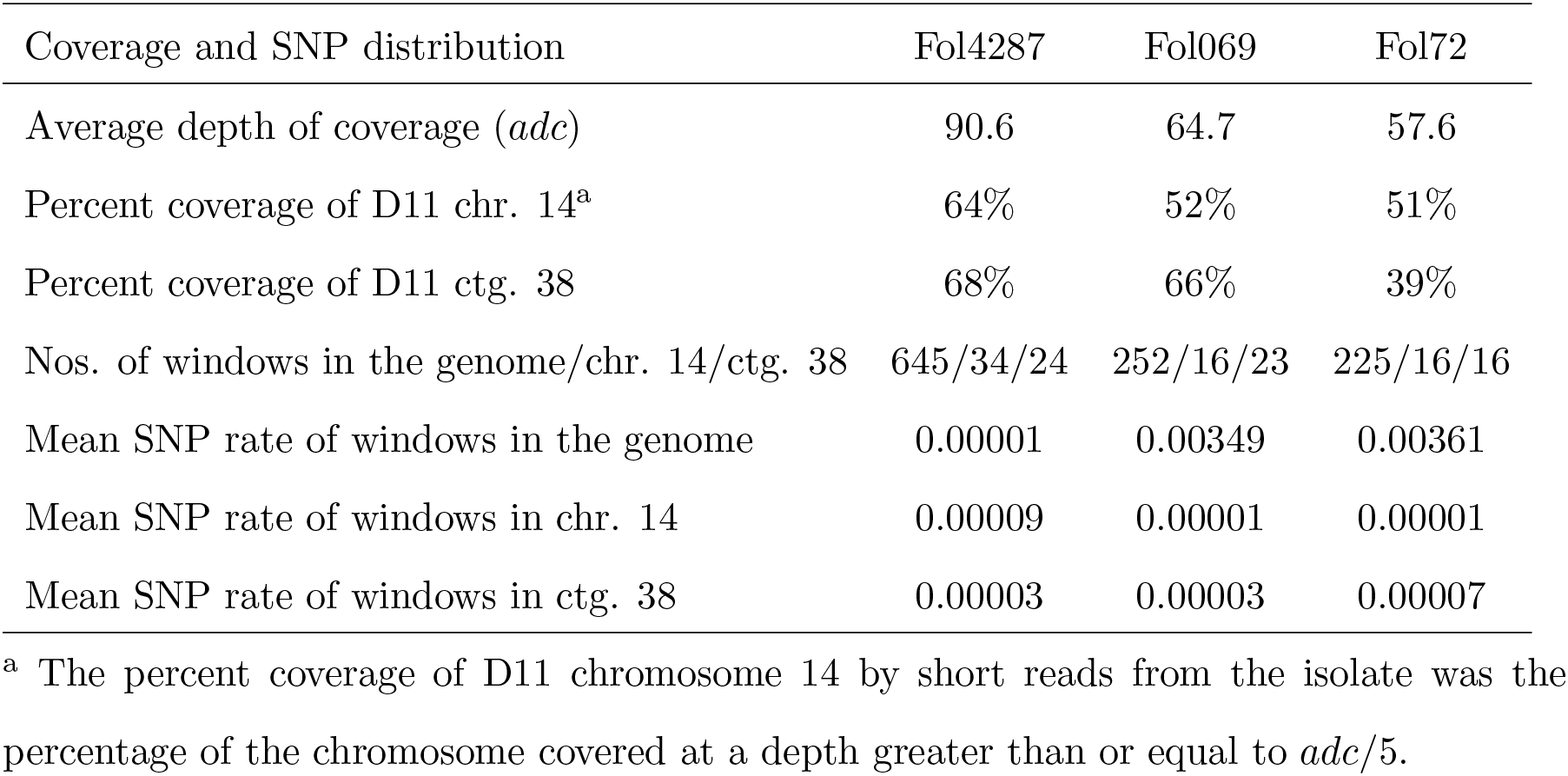
Mean SNP rates between Fol D11 and each of Fol4287, Fol069 and Fol072

| Coverage and SNP distribution | Fol4287 | Fol069 | Fol072 |
| --- | --- | --- | --- |
| Average depth of coverage ($adc$) | 90.6 | 64.7 | 57.6 |
| Percent coverage of D11 chr. 14<sup>a</sup> | 64% | 52% | 51% |
| Percent coverage of D11 ctg. 38 | 68% | 66% | 39% |
| Nos. of windows in the genome/chr. 14/ctg. 38 | 645/34/24 | 252/16/23 | 225/16/16 |
| Mean SNP rate of windows in the genome | 0.00001 | 0.00349 | 0.00361 |
| Mean SNP rate of windows in chr. 14 | 0.00009 | 0.00001 | 0.00001 |
| Mean SNP rate of windows in ctg. 38 | 0.00003 | 0.00003 | 0.00007 |

<sup>a</sup> The percent coverage of D11 chromosome 14 by short reads from the isolate was the percentage of the chromosome covered at a depth greater than or equal to $adc/5$ .

To see how unique the 3’ subtelomere of D11 chromosome 1 was to Fol, its sequence was compared to all genome assemblies (over 200) in the *Fusarium* genus. Of the top 35 matches, 19 were to the genome assembly of isolate D11, which were described above, and the 16 were at 99.66-99.99% identity over a length of 7.4-10.7 kb to genomic regions from other isolates in this *forma specialis*. The next 6 best matches were at 97-98% identity over 6.5-8.3 kb to isolates in other *formae speciales* of *F. oxysporum*.

Unlike subtelomeres in other *formae speciales* of *F. oxysporum* (see other subsections below), the D11 chromosome 1 subtelomere contained no genes encoding for proteins of more than 800 residues. Located upstream of the 3’ subtelomere of supernumerary chromosome 14 was a gene encoding for a protein of 2,102 residues with an OrsD domain (e-value = 1.5e-05). Immediately downstream of the 5’ subtelomere of chromosome 14 was an exon of 1,384 bp with an ORF encoding for DEAD (5.5e-11), Helicase_C (3.8e-10) and ResIII (6.7e-06) domains. Such a gene was found in the subtelomeres of core chromosomes in other *formae speciales* of *F. oxysporum* (see below). And chromosome 14 or contig 38 contained genes encoding for an Hsp70 protein (e-value = 1.3e-45), an MreB/Mbl protein (5.3e-12), and proteins with an Arrestin_N domain (3e-05), a bacterial SH3 domain (4.9e-05), an Est1 DNA/RNA binding domain (3.4e-12), a Cyclin_C domain (3e-09) and a Cyclin_N domain (1.5e-37), an MIS13 domain (1.5e-56), a Chromo domain (1.2e-08), a Chromo shadow domain (7.9e-17), a PARP domain (1.7e-28), a Homeodomain (1.5e-14), or an NDC10 II domain (2.1e-58). See Supplemental List for descriptions of domain name abbreviations.

### *F. oxysporum* f.sp. *radicis-cucumerinum*

In *F. oxysporum* f.sp. *radicis-cucumerinum* (Forc) isolate Forc016, its genome assembly consisted of 11 core chromosomes, one supernumerary chromosome (chromosome RC), and a number of unplaced contigs (van Dam et al. 2017). Chromosome RC, assembled from telomere to telomere, was flanked on either side by 13-kb complementary sequences at 99.96% complementarity (Fig. 1A). Of the core chromosomes, 7 were assembled from telomere to telomere and flanked on either side by 13-kb complementary sequences at 99.98-100% identity to and in the same orientation as those of chromosome RC (Fig. 1B), and the other 4 were flanked on one side by subtelomeres of this common type. Seven of the unplaced contigs were flanked on one side by common subtelomeres at 99.68-100% identity. Besides Forc016, this *forma specialis* includes isolates Forc024 and Forc031 (van Dam et al. 2016). The common Forc016 subtelomere was also present in multiple copies in these two isolates; its estimated copy number was 26 in isolates Forc024 and Forc031, and 29 in isolate Forc016.

The comparison of the common Forc016 subtelomere with the genome assemblies of all isolates in the *Fusarium* genus revealed that the subtelomere was at 100% identity over 12.9 kb to 6 *F. oxysporum* f.sp. melonis (Fom) isolates. Its estimated copy number was 2 or 3 in isolates Fom005, Fom006, Fom012, Fom013, and Fom016. Forc016 chromosome RC was much more conserved than its core chromosomes between Forc016 and these Fom isolates. For example, there were no SNPs between Forc016 and each of these isolates over a 31-kb region (starting at 766.8 kb) of Forc016 chromosome RC. In contrast, there were at least 60 SNPs over a 20.6-kb region (containing the conserved *EF-1α* gene) between Forc016 and each of these isolates. Note that both Forc and Fom isolates caused severe disease in musk melon (van Dam et al. 2016), a common host for both *formae speciales*. The Forc016 subtelomere was at 95-96% identity over 12.4 kb to 4 isolates from other *formae speciales*; it had no strong matches to other isolates.

**Figure 1.**
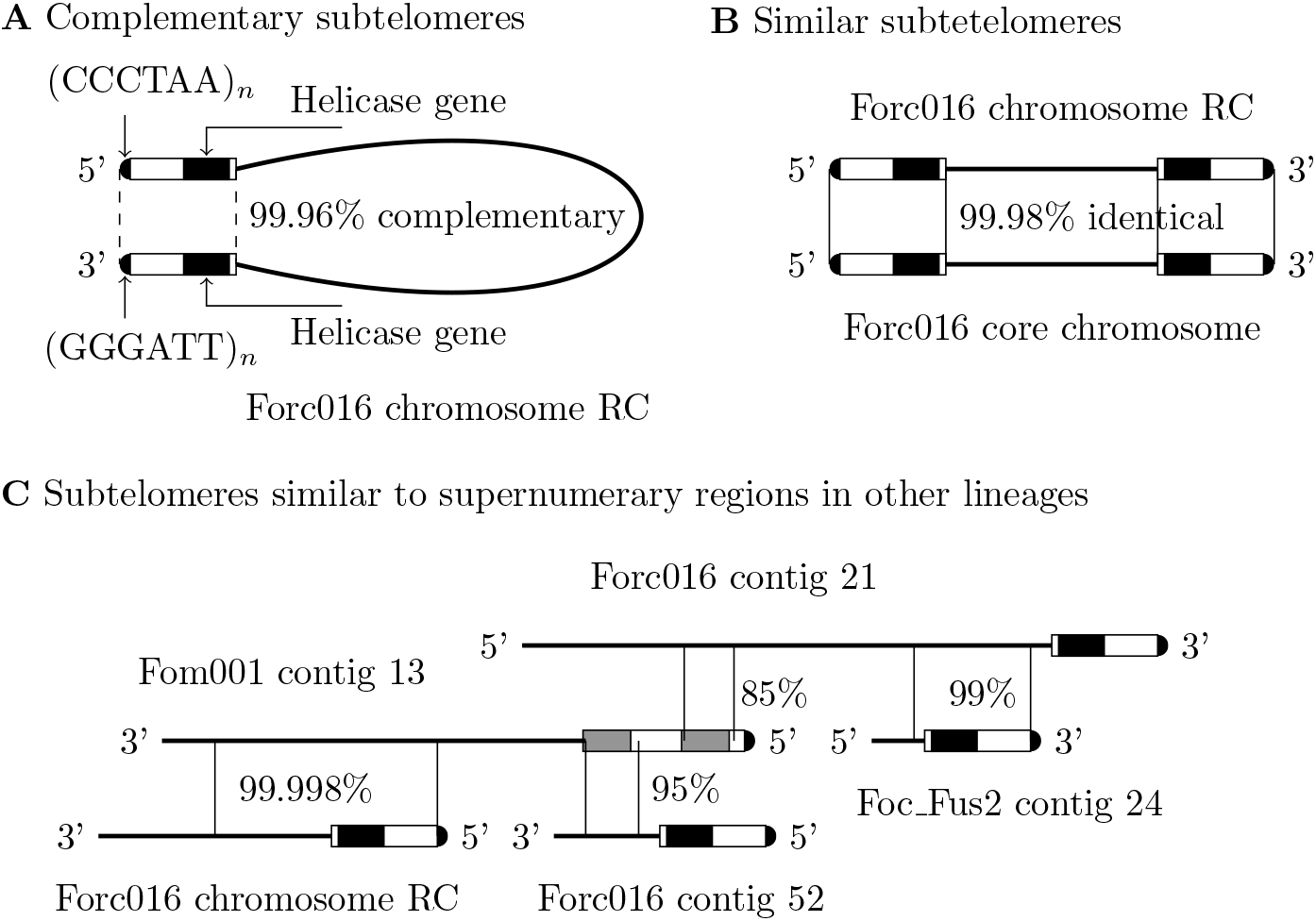
Three types of subtelomere sequence similarities. Each chromosome or contig is represented by a thick solid line with its name placed above or below the line. Similar regions between chromosomes/contigs are indicated by two vertical thin solid lines at their boundaries, with the percent identity given next to the lines. The subtelomere ends in a telomeric repeat (indicated by a half-filled circle) and contains a helicase gene (shown by a black rectangle). The figure is not drawn to scale. (*A*) The 5’ and 3’ subtelomeres of one single strand of Forc016 chromosome RC are complementary (indicated by two dashed lines at their boundaries) when read from both ends, making it possible for the supernumerary chromosome to repair its subtelomeres by forming a subtelomere hairpin if one of them is shortened. (*B*) The subtelomeres of core and supernumerary chromosomes at the same end are nearly identical, making it possible for them to exchange genes and repair subtelomeres by homologous recombination. (*C*) A subtelomere (in multiple copies) from one vegetative lineage is homologous to a non-terminal region (in a single copy) of a supernumerary chromosome or contig from another lineage. The subtelomere of Fom001 contig 13 contains a duplication (shown by a pair of gray rectangles), instead of a helicase gene.

In isolate Forc016, some chromosomes and contigs were also homologous over a length of 4 kb beyond the 13-kb subtelomere, with numerous C:G to T:A nucleotide changes indicative of RIP. For example, all next to their 3’ 13-kb subtelomeres, a region of chromosome 4 was at 90% identity to one of chromosome 10 with 394 C:G to T:A nucleotide changes out of a total of 396 substitutions, at 91% identity to one of contig 7 with 329 out of 332, and at 88% identity to one of contig 3 with 453 out of 456. An explanation for this observation is that RIP may have operated on the parts of these subtelomeres in an ancestral state.

And 5 of the unplaced contigs were not present in isolates Forc024 and Fore031. Two of these 5 contigs end in a telomeric repeat: the 3’ end of contig 55 and the 5’ end of contig 52. The 3’ subtelomere of contig 55 in forward orientation was at 100% identity over 7,100 bp to the 5’ one of contig 52 in reverse orientation, and the subtelomere contained a gene encoding for a protein of 1,531 residues with Helicase_C (e-value = 1.2e-09), DEAD (e-value = 3.2e-09) and ResIII (e-value = 5.3e-05) domains. This new subtelomere was not similar to those of the core chromosomes. And the longest of the 5 contigs, contig 53 of 987.8 kb, was at 95-98% identity over 36% of its length to supernumerary Fol D11 contig 38, and at 93-99% identity over 21% of its length to supernumerary Fol D11 chromosome 14. Thus, isolate Forc016 appeared to contain a supernumerary chromosome with a different host-specific subtelomere, which might have been acquired by horizontal transfer and maintained with the palindrome subtelomere at its ends.

The 3’ Forc016 subtelomere in supernumerary chromosome RC contained predicted genes encoding for a 393-residue protein with a kinase domain (e-value = 3.7e-25), and a 1,539-residue protein with Helicase_C (4.3e-12) and DEAD (3.6e-11) domains. The kinase and helicase genes were covered by RNA-seq reads (10 days post inoculation *in planta* conditions) to maximum depths of 78 and 28, respectively. The 5’ Forc016 subtelomere contained predicted genes encoding Helicase_C (5.9e-26), DEAD (6.7e-24), OrsD (7.2e-13), ResIII (8.6e-08). In addition to transposons and SM/pathogenicity genes, chromosome RC and contig 53 contained other types of genes encoding for proteins involved in DNA repair, recombination, cell cycle control, gene silencing in heterochromatin, chromosome segregation, chromatin architecture, responses to stress, and telomere elongation. See Supplemental List for a list of those genes.

### *F. oxysporum* f.sp *melonis*

In isolate Fom001 (van Dam et al. 2017), supernumerary contig 13 of 128.0 kb ends in a 5’ telomeric repeat. Contig 13 over positions 20.7 to 81.9 kb and over positions 81.8 to 111.8 kb displayed 99.99% identity to Forc016 chromosome RC over positions 0.1 to 61.3 kb and over positions 62.6 to 92.6 kb, respectively. The larger (with a length of 61 kb) of these two highly similar regions contained the 5’ subtelomere of chromosome RC (Fig. 1C), which was present at one or two ends of the 11 core chromosomes in isolate Forc016, but in a single copy (as an internal part of contig 13) in isolate Fom001. On the other hand, the 5’ subtelomere of contig 13 located at 1-19.3 kb, which was not present in isolate Forc016, was found at the telomere-containing ends of 17 contigs in isolate Fom001 at 98.69-99.76% identity for 6 of them over a length of 19 kb and for the rest over a length of 9.8-13.0 kb. Note that large portions of Forc016 chromosome RC were at 99.27-99.99% identity to parts of supernumerary contig 127 of 1,743,2 kb in isolate Fom001. Indeed, the mean SNP rates of windows in supernumerary contigs 13 and 127 were 26.63 and 20.19 times lower than that in the genome between isolates Fom001 and Forc016.

In isolate Fom001, the 5’ subtelomere of contig 13 was at 77% identity over positions 0.7 to 7.8 kb and at 63% identity over positions 10.4 to 20.7 kb to the 5’ subtelomere of contig 22 over positions 1.2 to 8.1 kb and over positions 1.2 to 9.8 kb, where the 63%-identity match included a block of 1,483 bp at 100% identity. This revealed a tandem duplication in the 5’ subtelomere of contig 13, where both copies were similar to parts of the 5’ subtelomere of contig 22. Moreover, the 5’ subtelomere of contig 13 was at 85% identity over one copy and at 95% identity over the other copy to a region of supernumerary Forc016 contig 21 and a region of supernumerary Forc016 contig 52, respectively (Fig. 1C). And contig 22 of 1,267,8 kb was supernumerary; it was enriched for transposons and SM/pathogenicity genes. The 5’ 9.8-kb subtelomere of contig 22 was at 99.27-99.96% identity to the 5’ subtelomeres of contigs 9, 15, 32, 86, 87, to a region of contig 107 from positions 9.9 to 19.7 kb, all in forward orientation, and to the 3’ subtelomere of contig 30 in reverse orientation.

Isolate Fom001 was more similar in core chromosome regions to the Fol lineage with isolate Fol007 than to other lineages including the Fom lineage (van Dam et al. 2016), and Fol D11 and Fom001 were at 99.97% identity over a 20.5-kb region containing the conserved *EF-1α* gene. However, the top matches between supernumerary Fom001 contig 22 and the *Fusarium* genome assemblies were all but one with isolates Fom004, Fom005, Fom006, Fom011, Fom012, Fom013 and Fom016 at 99.83-99.99% identity over several regions of 31.9 to 42.7 kb. More specifically, the mean SNP rate (0.00037) of windows in the Fom001 genome between isolates Fom001 and Fol007 was 9.38 times lower than that (0.00347) between isolates Fom001 and Fom006. On the other hand, supernumerary Fom001 contigs 13, 22 and 127 were not present in isolate Fol007. And between isolates Fom001 and Fom006, the mean SNP rates of windows in those three contigs were 23.93, 2.73 and 3.49 times lower than that in the Fom001 genome. The same observation was made for isolate Fom005.

The 5’ subtelomere of Fom001 contig 13 was covered over part 1 (1 bp to 9.8 kb) by short reads from some of the other Fom isolates; the portion from 13 to 15 kb was not covered by short reads from any of them. The estimated copy number of part 1 was 34 in Fom005, 27 in Fom006, 0 in Fom009, 44 in Fom010, 0 in Fom011, 28 in Fom012, 24 in Fom013 and 28 in Fom016; there was a problem with the Fom004 dataset. We found 1 to 3 SNPs in part 1 between Fom001 and each of the other 6 Fom isolates with multiple copies of part 1, and depths of low coverage from 1.3 to 1.6 kb for each isolate, suggesting variation over this region between Fom001 and the others. The estimated copy number of the 5’ subtelomere of Fom001 contig 22 was 0 in isolates Fom009, Fom010 and Fom011, and 4 to 7 in isolates Fom005, Fom006, Fom012, Fom013 and Fom016, with only 1 SNP at the same position between Fom001 and each of the other 5 isolates.

The 5’ 19-kb subtelomere of contig 13 contained 4 genes: two of them encoded for proteins with an MYND finger domain (4.1e-08 and 4.2e-10), and the other two encoded for hypothetical proteins of 1,340 and 1,255 residues. (In other words, each of the two copies in the 5’ subtelomere contained two genes, one encoding for an MYND finger domain and the other for a hypothetical protein.) About 8.7 kb downstream of the subtelomere was a gene (shared with Forc016 chromosome RC, see above) encoding for a 2,987-residue protein with DEAD (2.1e-28), Helicase_C (2.3e-27), ResIII (2.8e-06) domains. And the 5’ 9.8-kb subtelomere of contig 22 contained a gene encoding for a 403-residue protein with an MYND finger domain (1.3e-09) and another gene encoding for a hypothetical protein of 1,543 residues. About 4.5 kb downstream of the subtelomere was a gene encoding for a 1,317-residue protein with DEAD (1.2e-10), Helicase_C (1.5e-07), and DUF3505 (6.2e-05) domains. And immediately downstream of this gene was another gene encoding for a 2,649-residue protein with DUF3505 (3.5e-13) and DEAD (2.5e-06) domains.

### *F. oxysporum* f.sp. *cubense*

*F. oxysporum* f.sp. *cubense* (Focb) is a *forma specialis* of polyphyletic lineages causing Panama disease of banana (O’Donnell et al. 1998). One group of lineages is Focb race 1, which attacks a banana cultivar named ‘Gros Michel’ and caused the 20th century epidemic; another group is Focb tropical race 4 (TR4), which affects a banana cultivar named ‘Cavendish’ (resistant to race 1) and the hosts of race 1 (Zheng et al. 2018). For Focb race 1 isolate 160527, a high-quality genome assembly was recently generated (Asai et al. 2019); for Focb TR4 isolate II5, a draft genome assembly was generated years ago (GenBank accession: GCA 000260195.2). Focb 160527 had a total of 15.7 Mb strong matches (of length above 1 kb and at identity above 98.7%) to isolate Forc016 (not in Focb), as compared to a total of 6.9 Mb strong matches to Focb TR4 II5, suggesting that Focb 160527 was more closely related to Forc016 than to Focb TR4 isolate II5, which is consistent with the polyphyletic characterization of Focb.

The Focb 160527 assembly consisted of 12 contigs with 20 terminal regions ending in telomeric repeats. One of the contigs, contig 2 of 5,885.8 kb with each terminal region ending in a telomeric repeat, was a supernumerary chromosome flanked on each side by complementary subtelomeres of 8.5 kb at 99.98% complementarity. Note that this supernumerary chromosome was longer than all but one contig, with 7 of them assembled from telomere to telomere. The 3’ subtelomere of supernumerary contig 2 was at 98.83-99.98% identity to 20 contig terminal regions in forward or reverse orientation. Another contig, contig 12 of 1,261.1 kb, was also supernumerary with th same kind of complementary subtelomeres as contig 2. Note that no match over the Focb 160527 supernumerary subtelomere was found between the assemblies of Focb 160527 and Focb TR4 II5.

To see if part of the 3’ Focb 160527 supernumerary subtelomere was present in Focb TR4 isolates, short reads from three Focb TR4 isolates La-2, My-1 and Vn-2 (Zheng et al. 2018) were mapped onto the subtelomere as a reference. A 1.6-kb region of the subtelomere at a distance of 3.2 kb from its 3’ end was covered at a maximum depth of 538 for La-2, 1,587 for My-1, and 1,988 for Vn-2. An estimated copy number for this region was 24 in La-2, 20 in My-1, 22 in Vn-2. A total of 14 SNPs between the reference and short reads, with each SNP shared by the three TR4 isolates, were found in this region. The Focb 160527 subtelomere contained a gene encoding for a 1454-residue protein with a DEAD domain (1.3e-08), a Helicase_C domain (1.7e-08) and an OrsD domain (9.7e-05). The region of the subtelomere covered by short reads from the three TR4 isolates included all of the Helicase_C domain and the C-terminal half of the DEAD domain. Although Focb 160527 and Focb TR4 II5 were not closely related, the three TR4 isolates contained a region in 20-24 copies that was at 99.1% identity to a region of the Focb 160527 subtelomere. And the Focb 160527 assembly was used as a reference for mapping short reads from the three TR4 isolates, more than 64% percent of contig 2 (or 96% of contig 12) were not covered by short reads from each of the three TR4 isolates; about 18-58% percent of every other contig were not covered by short reads from each of the three TR4 isolates. Although Focb TR4 isolates had little variation with low SNP rates (less than 0.00001) between the Focb TR4 II5 reference and short reads from other TR4 isolates (Zheng et al. 2018), two regions of Focb 160527 contig 2 were found with little variation with two of the three TR4 isolates but with significant variation with the other isolate. One was a 29.6-kb region starting at 3,645.9 kb of contig 2 that was covered with no SNPs at average depths of 56.4 (for My-1) and 55.2 (for Vn-2) but at an average depth of 0.1 for La-2. The other was a 47.4-kb region starting at 4,006.6 kb of contig 2 with one common SNP at average depths of 18.3 (for La-2) and 54.8 (for Vn-2) but at an average depth of 0.0 for My-1.

Besides transposons, supernumerary contig 2 contained genes encoding for 2 proteins each with a Cyclin_N domain (6.4e-26 and 4.4e-07), 4 proteins each with a PARP/PARP_reg domain (3.9e-32, 1.6e-20, 1.9e-20 and 2.7e-22), an HSP70 protein (1e-57), a protein with RCC1 and RCC1_2 domains (2.4e-66 and 5.6e-40), 3 proteins each with a Lactamase_B domain 2.1e-06, 4.9e-11 and 6.4e-17), and 2 proteins each with a Lysm domain (3e-06 and 4.3e-15).

### *F. oxysporum* f.sp. *cepae*

In *F. oxysporum f.sp. cepae* (Foc) isolate Foc_Fus2, its genome assembly consisted of 46.7-Mb core chromosome regions and 5.7-Mb supernumerary regions (Armitage et al. 2018). Chromosome 7 and contig 25 end in 5’ telomeric repeats, and contig 24 and chromosomes 2 and 12 end in 3’ telomeric repeats. The 3’ subtelomeres of the last three contigs/chromosomes in forward orientation were at 98.89-99.97% identity (over a length of 9.3 kb) to the 5’ subtelomeres of the first two in reverse orientation. And the 3’ subtelomere of contig 24 had more similarities (at 99.53-99.98% identity over 4.5-7.6 kb) with the ends of 5 contigs and 1 chromosome that do not end in a telomeric repeat. To shed light on its origin, this subtelomere was found to be highly similar to genomic regions of isolates from 8 other *formae speciales* of *F. oxysporum*. For example, the top match was at 99.75% identity over a length of 6.4 kb with contig 21 of 73 kb from isolate Forc016, and the next one was at 99.74% over a length of 6.3 kb with contig 533 of 6.3 kb from isolate Fol069. The conservation levels of these two matches were close to those (at 99.83% and 99.66% identity) of the matches between isolate Foc_Fus2 and each of isolates Forc016 and Fol069 over the *EF-1α* gene. While Fol069 contig 533 was short, Forc016 contig 21 ends in a 3’ telomeric repeat, next to which was a copy of the 13-kb Forc016 subtelomere. Located 6 kb upstream of this subtelomere was the region of Forc016 contig 21 that was similar to the Foc_Fus2 subtelomere (Fig. 1C); this region encoded for a protein of 1,127 residues with a DEAD domain (e-value = 7e-09). Note that Forc016 contig 21 was supernumerary.

We also checked whether the Foc_Fus2 subtelomere was present in multiple copies in other isolates from its *forma specialis*. We found two more isolates, A23 and 125, for which and for Foc_Fus2, genomic data sets of short reads are available (Armitage et al. 2018). The estimated copy number of the subtelomere was 26 in isolate Foc_Fus2, 30 in isolate A23, and 25 in isolate 125. Note that the 200-bp region immediately upstream of the subtelomere was covered only by reads from Foc_Fus2, and a 5-bp region inside the subtelomere was covered 10 times more by reads from Foc_Fus2 and 125 than from A23. This suggests variation among the isolates in genomic regions containing copies of the subtelomere. And the subtelomere in contig 24 contained a gene encoding for a protein of 1411 residues with Helicase_C (e-value = 9.1e-09) and DEAD (1.1e-08) domains, and the rest of contig 24 contained transposons and pathogenicity genes.

### F. equiseti

A genome assembly (Accession: GCA 003313175.1) of *F. equiseti* isolate D25-1 was composed of 16 contigs, 5 of which end in 3’ telomeric repeats. These 5 contigs in the same orientation were highly homologous (at 99.63-99.73% identity) over a 3’ subtelomeric region of 10.4 to 10.7 kb. Two of these contigs, scaffold 1 of 8,345 kb and scaffold 6 of 4,117 kb, were parts of core chromosomes, where the regions of the scaffolds conserved at ≥ 94% identity with *F. equiseti* isolate CS5819 were covered by short reads from this isolate. (The pathogenicity status of this isolate is unknown). Another two of them, scaffold 8 of 556 kb and scaffold 11 of 183 kb, were supernumerary, since they contained no homologs of essential genes except ones encoding for DNA pol A 3’-5’ exonuclease (e-value = 4.3e-09), Centromere protein Scm3 (e-value = 4.7e-15), and a Mif2/CENP-C protein (e-value = 3e-17), which were not conserved with isolate CS5819. The last one, scaffold 15 of 10.5 kb, was too short to determine. The D25-1 subtelomere was not covered by short reads from isolate CS5819. The subtelomere contained a gene encoding for a hypothetical protein of 1,830 residues and another one for a protein of 402 residues with a MYND finger domain (e-value = 1.6e-09). And the subtelomere was at 92-96% identity to genomic regions mostly from *F. oxysporum* isolates; for example, one of these regions was from supernumerary chromosome 14 of isolate Fol4287.

### Colletotrichum higginsianum

*Colletotrichum higginsianum* isolate IMI 349063 contained 12 chromosomes assembled from telomere to telomere (except chromosome 7, consisting of more than one contig) (Dallery et al. 2017). Two of them were supernumerary: chromosomes 11 and 12 of 646 and 598 kb, respectively; the rest were core with a size of 3 to 6 Mb (Dallery et al. 2017). The 3’ ends (in forward orientation) and 5’ ends (in reverse orientation) of the 12 chromosomes were highly homologous and were classified into three types of sequences (Dallery et al. 2017). Here we examined the levels of diversity in these sequences to shed light on their origin in connection with the supernumerary chromosomes. Chromosome 12 was flanked on either side by nearly complementary sequences of 12,772 and 11,195 bp; an alignment of the sequences in opposite orientation contained two sections of similar sequences (at 98.14% identity) that were separated by a section of different sequences of 1,926 and 341 bp (at overall 83% identity over the three sections). This subtelomeric element is called a 11-kb element, based on the number of exact base matches (10,653) in the alignment. And chromosome 11 was similarly associated with 18-kb complementary sequences at 99.87% complementarity. Except for a 7.6-kb insertion and a 1.9-kb deletion (or a 0.3-kb deletion), the 18-kb element was 99.01% (or 98.14%) identical to the 5’ (or 3’) copy of the 11-kb element. As for the other chromosomes, their ends were at 99.60-99.99% identity to one of the ends of chromosome 12 over the 11-kb element, except for the 5’ ends of chromosomes 7 and 9, which were at 99.91-99.88% identity to the 3’ end of chromosome 11 over the 18-kb element.

The data suggest that all the duplications of the 18-kb element were more recent than some of the duplications of the 11-kb element. The analysis found supernumerary chromosome 11 with a copy of the 18-kb element on either end, but no core chromosome with such an arrangement, just two core chromosomes (chromosomes 7 and 9) with a copy only on the 5’ end. This sheds light on the question of whether the supernumerary chromosomes with complementary subtelomeric sequences arose before the core chromosomes acquired the same type of sequences on both of their ends.

Moreover, between *C. higginsianum* isolates IMI 349063 and MAFF 305635, chromosome 12 was more variable than chromosome 11 and the core chromosomes, which were at similar levels of conservation (Plaumann et al. 2018). This species illustrates a case where a pathogenic supernumerary chromosome (chromosome 11) was less variable than a non- or less-pathogenic supernumerary chromosome (chromosome 12) between the isolates (Plaumann et al. 2018). The estimated copy number of the 3’ subtelomeric element of chromosome 12 in *C. higginsianum* isolate MAFF 305635 was 22. A 1.6-kb region of the 3’ subtelomeric element of chromosome 11 was not covered by short reads from this isolate, suggesting that the element differed from those in the isolate.

Chromosome 12 contained genes encoding for a 477-residue protein from DNA polymerase family B (e-value = 7.4e-34), for a 336-residue protein with domain 2 of RNA polymerase Rpb1 (2.5e-08), for a 244-residue protein with domain 6 of RNA polymerase Rpb2 (2.7e-13), and for a 182-residue protein with a chromo (CHRromatin Organization MOdifier) domain (3.4e-23), as well as many transposons and pathogenicity genes. Some of the genes on chromosome 11 encoded for a 384-residue protein from glycosyl hydrolases family 18 (1.7e-79), a 499-residue protein with a HET domain (2.6e-13), and a 1,276-residue protein with a Helicase_C domain (1.2e-05) and an SNF2 family N-terminal domain (5.3e-05). Besides predicted helicase domains (Dallery et al. 2017), the protein encoded by the gene in the 18-kb element included the following domains: chromo (e-value = 1.4e-15), OrsD (1.2e-11), and zinc knuckle (2.5e-06).

### Pyrenochaeta lycopersici

*Pyrenochaeta lycopersici* isolate CRA-PAV ER 1518 was used to study the evolution of diverse subtelomeres. A genome assembly of this isolate included 13 contigs ending in 3’ telomeric repeats. These contigs contained no highly similar matches over their 3’ sub-telomeres. On the other hand, some of these 3’ subtelomeres were similar to other regions of the assembly. For example, consider three of the 13 contigs: contig 205 of 23.8 kb, contig 206 of 85.7 kb, and contig 53 of 274.7 kb. Except for a 5’ end of 1,308 bp and a 3’ end of 104 bp, contig 205 was at 99.89% identity over a length of 22.4 kb to a 5’ end of contig 206, both in forward orientation. A 3’ end (excluding the last 730 bp) of contig 53 was at 90.62% identity over a length of 12.6 kb to a 5’ end (excluding the first 892 bp) of contig 205, both in forward orientation. The alignment of the two contig ends contained a total of 993 substitutions, of which, 957 (96.4%) were C:G to T:A nucleotide changes indicative of RIP, with 936 (97.8%) of those changes involving A’s or T’s in the 5’ end of contig 205. This end of contig 205 contained no long ORFs. But downstream of it, contig 205 over its 3’ subtelomere contained a gene encoding a 1,439-residue protein with several domains: OrsD (e-value = 7e-23), Helicase_C (4.8e-11), DEAD (1.2e-10), and ResIII (7.9e-05). The 3’ subtelomere of contig 53 contained a gene encoding a 1,463-residue protein with an OrsD domain (3.9e-10). Except for its 5’ end, contig 206 contained no significant matches to other contigs, and no long ORFs were found in its 3’ terminal region. As more examples of matches with contig 205, portions of its 5’ AT-rich region were at 99.66-99.96% identity over a length of 9.8-12.5 kb to 9 other regions, and portions of its 3’ helicase-encoding region were at 99.23-99.39% identity over a length of 10.2 kb to 4 other regions. There were many other significant matches involving some of the 3’ subtelomeres (details omitted). Highly similar matches involving the 3’ subtelomeres with long ORFs indicate that their duplications were recent and that RIP might have been involved in generating some of their AT-rich copies.

### HET domain genes

We collected data on the number, location and variation of HET domain genes in isolate Fol D11 to shed light on their origin and role. We predicted 135 HET domain genes with an e-value below 1.e-05 in the Fol D11 genome assembly. Of these genes, 23 (17%) were located in the supernumerary chromosomes or contigs, where a contig was classified as being supernumerary if its coverage by short reads from isolates Fol069 and Fol072 was less than 60% and it was enriched for transposons and SM/pathogenicity genes. [At least 73% (or 74%) of each D11 core chromosome was covered by short reads from Fol069 (or Fol072), and 56.4% (or 55.1%) of supernumerary Fol D11 chromosome 14 was covered by short reads from Fol069 (or Fol072).] Of the remaining 112 HET domain genes, 91 were found in the 9 core chromosomes and 21 in core contigs 14 and 16.

We evaluated the variation between D11 and each of isolates Fol4287, Fol069, Fol072 and Fo47 by examining the coverage of the D11 genomic regions around each of the 91 HET domain gene loci by short reads from each of these 4 isolates. Here the genomic region around a gene was defined as an area centered in the gene and of a length twice that of the coding region. Except for a 3’ 500-kb end region of chromosome 1 with 3 HET domain genes and a 5’ 40-kb end region of chromosome 11 with 1 HET domain gene, the D11 genomic regions around the loci of the HET domain genes were covered at sufficient depths (≥ 10) without any variation by short reads from Fol4287. However, some positions in each of the D11 genomic regions around 89 of the 91 HET domain genes were not covered by short reads from Fol069, Fol072 or Fo47. This shows that the core genomic regions around the HET domain genes tended to be free of variation between isolates D11 and Fol4287 from the same lineage, but were variable between D11 and each of isolates Fol069, Fol072 and Fo47 in different lineages from D11. The 91 HET domain genes were located in regions with SM/pathogenicity genes; for example, 40 of the 91 genes were located within 15 kb of a gene encoding for a major facilitator superfamily (MFS) or cytochrome P450 protein. Similar observations were made about the 21 HET domain genes in core contigs 14 and 16; for example, 16 of these genes were located within 15 kb of genes encoding for MFS or P450 proteins.

## Discussion

In several asexual species of filamentous fungal pathogens with supernumerary chromosomes, supernumerary and core chromosomes are flanked on either side by highly complementary long sequences. This subtelomeric sequence often contains the open reading frame of a helicase gene, like the Y’ element in the budding yeast, whose helicase is expressed many fold in the absence of telomerase (Yamada et al. 1998). Thus, this subtelomeric homology structure may have a role in maintaining telomeres for supernumerary chromosomes as well as core ones during vegetative reproduction. The structure also suggests homologous recombination as a mechanism for the exchange of genes between supernumerary chromosomes and the ends of core chromosomes, explaining the H3K27me3 connection between the ends (large subtelomeric blocks of non-syntenic DNA) and the odds (supernumerary chromosomes) (Galazka and Freitag 2014). Moreover, supernumerary chromosomes are enriched for genes involved in modulating chromatin structure. And a large number of HET domain genes preserved within isolates from the same species or *forma specialis* may have a role in maintaining their neighboring non-essential SM/pathogenicity genes, because a loss of a DNA segment with a HET domain gene may result in cell death. In Fol, some isolates are variable around most HET domain genes but are more conserved in core and supernumerary subtelomeres as well as in whole supernumerary chromosomes. This suggests that the subtelomeres and supernumerary chromosomes are host-specific. Thus, the subtelomeric homology structure on the core and supernumerary chromosomes in several asexual species may be used as genomic signatures of host adaptation or host-switching.

In Fol and between Forc and Fom, a supernumerary chromosome or parts of it are more conserved than the core chromosomes. This, combined with the *in vitro* experimental evidence of HCT with the supernumerary chromosome (Ma et al. 2010 and van Dam et al. 2017), suggests that HCT of supernumerary chromosomes between different asexual lineages occurs in nature. These asexual lineages are formed to acquire and maintain supernumerary chromosomes through the duplication of subtelomeres at the ends of core chromosomes. The highly conserved subtelomeric structure also suggests that HCT occurs recently. The presence of a new subtelomere with a long insertion in some core and supernumerary chromosomes indicates that the subtelomeric homology structure evolves as an ongoing process to acquire and maintain the most-pathogenic supernumerary chromosome in the *forma specialis*. Consequently, HCT occurs frequently in asexual filamentous fungal pathogens with supernumerary chromosomes. Supernumerary chromosomes function as a powerful tool of evolution for non-essential but beneficial genes in filamentous fungi: they carry and evolve such genes across a wider species complex of filamentous fungi in a complementary mechanism to the Mendelian process. Some of these genes are homologs of essential genes, involved in DNA replication and repair, and gene expression and chromosome homeostasis. Other regions such as their complementary subtelomeres come from supernumerary chromosomes in other lineages. Taken together, HET domain genes along with SM/pathogenicity genes are acquired by HCT of supernumerary chromosomes, and exchanged between core and supernumerary chromosomes by homologous recombination over subtelomeres.

Core chromosomes contained numerous C:G to T:A nucleotide changes indicative of RIP, while supernumerary chromosome did not, which is consistent with the study by Vanheule et al. (2016). And no such changes were observed in the highly similar subtelomeric sequences of core chromosomes. An explanation consistent with these observations is given as follows. A filamentous fungal pathogen alternates between sexual and asexual cycles (Nieuwenhuis and James 2016). During the sexual cycle, highly similar regions in its genome, like highly homologous subtelomeric regions, sustain mutations by processes like RIP. The pathogen evolves by genome wide homologous recombination within the population. During the asexual cycle, the pathogen first acquires (by HCT) supernumerary chromosomes with highly complementary subtelomeric sequences, which are subsequently duplicated at its core chromosomes. Then the pathogen evolves by localized homologous recombination over subtelomeres and by HCT of supernumerary chromosomes within and between populations. The asexual cycle allows some homologs of essential genes as well as SM/pathogenicity genes to evolve through gene duplication and HCT, without being suppressed by processes like RIP. The asexual cycle could last for thousands of generations; it was estimated that the wild yeast *Saccharomyces paradoxus* had an asexual cycle of 1,000 generations (Tsai et al. 2008). Each asexual cycle corresponds with the emergence of a new asexual lineage. So the length of the asexual cycle may be related to the frequency of speciation events.

An explanation for the complementary long sequences with a helicase gene at the ends of a supernumerary chromosome is that the helicase and long complementary sequence structure are involved in maintaining the stability of the chromosome by lengthening its telomeres based on homologous recombination in vegetative cells. This arrangement is especially important when the supernumerary chromosome is transferred horizontally into a different species whose chromosome ends are not homologous to those of the supernumerary chromosome.

Certain asexual filamentous fungal pathogens adapt to their hosts by acquiring supernumerary pathogenicity chromosomes horizontally and duplicating the supernumerary subtelomeres at the ends of their core chromosomes. An evolution-guided strategy for controlling these pathogens is to develop a comprehensive identification method based on their host-specific subtelomere structures and to deploy the method globally to identify and prevent them from disseminating their supernumerary pathogenicity chromosomes.

## Methods

We obtained the genome assemblies of the following isolates (by their GenBank assembly accessions) from GenBank at National Center for Biotechnology Information (NCBI): *C. higginsianum* isolate IMI 349063 (GCF_001672515.1), *F. equiseti* isolate D25-1 (GCA_003313175.1), *F. oxysporum* f.sp. *cepae* isolate Fus2 (GCA_003615085.1), *F. oxysporum* f.sp. *cubense* race 1 isolate 160527 (GCA 005930515.1), *F. oxysporum* f.sp. *cubense* tropical race 4 isolate II5 (GCA_000260195.2), *F. oxysporum* f.sp. *lycopersici* race 3 isolate D11 (GCA 003977725.1), *F. oxysporum* f.sp. *melonis* isolate Fom001 (GCA_002318975.1), *F. oxysporum* f.sp. radicis-cucumerinum isolate Forc016 (GCA_001702695.2).

Pairwise comparisons of genome assemblies were performed with MUMmer3 (Kurtz et al. 2004) and DDS2 (Huang et al. 2004). Comparisons of a genome assembly with a database of genome assemblies were made with Blastn (Altschul et al. 1990). An alignment of two genomic regions with sections of similar regions separated by sections of different regions was computed with GAP3 (Huang and Chao 2003). Multiple local alignments between two sequences were computed with SIM (Huang and Miller 1991). Genes in a genome assembly were identified with Augustus (Stanke and Waack 2003) and AAT (Huang et al. 1997). Domains in protein sequences were found with HMMER (Finn 2011).

We also obtained the datasets of short reads for the following isolates (by their SRA accessions) from Sequence Read Archive (SRA) at NCBI: *C. higginsianum* isolate MAFF 305635 (SRR6412364); *F. equiseti* isolate CS5819 (SRR5194938); *F. oxysporum* isolate Fo47 (SRR306682, SRR306671, SRR306668, SRR306667, SRR306657); *F. oxysporum* f.sp. *cepae* isolates 125 (SRR4408417), A23 (SRR4408418), Fus2 (SRR4408416); *F. oxysporum* f.sp. *cubense* tropical race 4 isolates La-2 (SRR7226878), My-1 (SRR7226877), Vn-2 (SRR7226879); *F. oxysporum* f.sp. *lycopersici* isolates Fol007 (SRR3142257, SRR3142258), Fol014 (SRR307256, SRR307271), Fol026 (SRR307236, SRR307255, SRR307293), Fol069 (SRR307106, SRR307107, SRR307113, SRR307123, SRR307257, SRR307115, SRR307266), Fol072 (SRR307122, SRR307092, SRR307091, SRR307090, SRR307086, SRR307281, SRR307250), Fol4287 (SRR3139043, SRR7690004); *F. oxysporum* f.sp. *melonis* isolates Fom004 (SRR1343458), Fom005 (SRR1343500), Fom006 (SRR1343569), Fom012 (SRR1343574), Fom013 (SRR1343575), Fom016 (SRR1343577); *F. oxysporum* f.sp. radicis-cucumerinum isolates Forc016 (SRR3139027, SRR3139028), Forc016 *in planta* RNA-seq (SRR5666309), Forc024 (SRR3139029, SRR3139030), Forc031 (SRR3139031, SRR3139032).

Isolate Fo47 is non-pathogenic, the pathogenicity status of isolate CS5819 is unknown, and all the other isolates are pathogenic.

Short reads were mapped onto a genome assembly as a reference, as previously described (Huang 2014; Huang et al. 2016). Note that the highly similar subtelomeres of core and supernumerary chromosomes in the genome assembly were not covered by any short reads, because no short reads can be uniquely mapped to any duplicated regions in the reference. To determine whether a subtelomere was present in the genome represented by short reads, the sequence of the subtelomere was used alone as a reference for mapping the short reads. To estimate the copy number of the subtelomere in the genome represented by short reads, the average coverage depth of the subtelomere by short reads was calculated, and also calculated (for comparison) was that of a genomic region containing the *EF-1α* gene. The estimated copy number of the subtelomere was defined as the average coverage depth for the subtelomere divided by that for the *EF-1α* gene region. Note that the coverage depth at each base position of a genomic region was calculated by using Bedtools with the genomecov subcommand (Quinlan and Hall 2010).

## Data Access

No new data were generated in this project.

## Acknowledgments

This research was conducted during X.H.’s Faculty Professional Development Assignment in 2018-2019.

## Supplemental List

### Abbreviations for protein domains

- BRCT, BRCA1 C Terminus (BRCT) domain;
- Chitin_bind_1, Chitin recognition protein;
- CHAT, CHAT domain;
- CinA, Competence-damaged protein;
- Clr5, Clr5 domain;
- Cyclin_N, Cyclin N-terminal domain;
- Cyclin_C, Cyclin C-terminal domain;
- DDE_1, DDE superfamily endonuclease;
- DEAD, DEAD/DEAH box helicase;
- DnaJ, DnaJ domain;
- DNA_pol_A_exo1, 3’-5’ exonuclease;
- Dynamin_N, Dynamin family;
- Dynamin_M, Dynamin central region;
- EST1_DNA_bind, Est1 DNA/RNA binding domain;
- F-box, F-box domain;
- FTA2, Kinetochore Sim4 complex subunit FTA2;
- GCR1_C, a transcriptional activator of glycolytic enzymes;
- Glyco_hydro_18, Glycosyl hydrolases family 18;
- Helicase_C, Helicase conserved C-terminal domain;
- Helo_like_N, Fungal N-terminal domain of STAND proteins;
- HeLo, Prion-inhibition and propagation;
- HlyIII, Haemolysin-III related;
- Lactamase_B, Metallo-beta-lactamase superfamily;
- MIS13, Mis12-Mtw1 protein family;
- Myb_DNAbinding, Myb-like DNA-binding domain;
- Na_Ca_ex, Sodium/calcium exchanger protein;
- NDC10_II, Centromere DNA-binding protein complex CBF3 subunit;
- OrsD, Orsellinic acid/F9775 biosynthesis cluster protein;
- PARP, Poly(ADP-ribose) polymerase catalytic domain;
- PARP_reg, Poly(ADP-ribose) polymerase, regulatory domain;
- PHM7_cyt, Cytosolic domain of 10TM putative phosphate transporter;
- PHM7_ext, Extracellular tail of 10TM putative phosphat transporter;
- RCC1, Regulator of chromosome condensation (RCC1);
- RCC1_2, Regulator of chromosome condensation (RCC1);
- ResIII, Type III restriction enzyme, res subunit;
- RSN1_7TM, Calcium-dependent channel, 7TM region;
- RSN1_TM, Late exocytosis, associated with Golgi transport;
- Scm3, Centromere protein Scm3;
- SET, SET domain;
- WGR, WGR domain;
- VWA, von Willebrand factor type A domain.

### Genes in Forc016 supernumerary chromosome RC

- a 291-residue protein with Cyclin_N (5.2e-26) and Cyclin C (1.3e-07) domains,
- a 241-residue protein with a Cyclin_N domain (4.4e-07),
- 2 392-residue proteins with Cyclin_N (3.4e-39) and Cyclin C (4.7e-10) domains,
- a 164-residue protein with a chromo domain (4.5e-13) and a chromo shadow domain (1.3e-05),
- a 301-residue protein with a chromo shadow domain (2.6e-18) and a chromo domain (2e-11),
- a 791-residue protein with NDC10_II (5.8e-59) and GCR1_C (5.5e-19),
- a 1112-residue protein with NDC10_II (1.2e-58), DDE_1 (1.4e-26), and GCR1_C (7.7e-19),
- 2 785-residue proteins with NDC10_II (1.6e-49) and GCR1_C (3.6e-19),
- a 1174-residue protein with NDC10_II (4.8e-49), DDE_1 (1.4e-26), and GCR1_C (8.1e-19),
- a 429-residue protein with a haemolysin-III related domain (2.9e-37),
- a 601-residue protein with PARP (8.3e-28), PARP_reg (4.4e-09), WGR (6.5e-07), and BRCT (1.4e-05) domains,
- a 656-residue protein with PARP (4.2e-49), PARP_reg (1.4e-42), WGR (9.5e-22), and BRCT (3.1e-07) domains,
- 2 439-residue proteins with PARP (1.2e-38), PARP_reg (5.2e-20) domains,
- a 561-residue protein with PARP (4.7e-24), BRCT (3.5e-08), and WGR (1.3e-06) domains,
- a 557-residue protein with PARP (1.6e-20), and WGR (2.4e-16) domains,
- 2 432-residue protein with a Glyco_hydro_18 domain (7e-75),
- a 366-residue protein with a Glyco_hydro_18 domain (2.5e-71),
- a 392-residue protein with a Glyco_hydro_18 (4.9e-47), and Chitin bind 1 (7.9e-05) domains,
- 3 326-residue proteins with a Glyco_hydro_18 domain (3.4e-70) domain,
- a 305-residue protein with a WGR domain (7e-09),
- a 246-residue protein with a WGR domain (4e-12),
- a 220-residue protein with a CHAT domain (1.9e-07),
- a 301-residue protein with a VWA domain (6.9e-08),
- a 872-residue protein with a RSN1_7TM domain (3.6e-64),
- a 419-residue protein with a DnaJ domain (1.5e-22),
- 2 517-residue proteins with a EST1_DNA bind (2.9e-13) domain,
- a 247-residue protein with a RSN1_7TM domain (1e-16),
- a 2850-residue protein with a Na_Ca_ex domain (3.5e-37),
- a 480-residue protein with RCC1 (1.9e-66) and RCC1 2 (5.5e-40) domains,
- a 369-residue protein with a SET domain (1.1e-09),
- a 621-residue protein with a Clr5 domain (5.9e-05),
- a 221-residue protein with a F-box domain domain (9.1e-05),
- a 264-residue protein with a DNA_pol_A exo1 domain (3.2e-08),
- a 460-residue protein with a MIS13 domain (3.3e-43),
- a 892-residue protein with RSN1_7TM (4.5e-75), RSN1_TM (1.5e-41), PHM7_cyt (5.8e-34), PHM7_ext (1.1e-24) domains.

### Alternative genes in Forc016 supernumerary contig 53

- A 341-residue protein with a Forkhead domain (2e-16),
- a 829-residue protein with 4 PDZ domains (4.1e-08 to 8.9e-18),
- a 440-residue protein with 2 Myb_DNA-binding domains (5.4e-06 and 6.6e-10),
- a 179-residue protein with a BTB/POZ domain (2.9e-06),
- a 535-residue protein with Hsp70 (8.4e-41) and MreB/Mbl (7e-08) domains,
- a 765-residue protein with a Helo_like_N domain (3.1e-07),
- a 787-residue protein with FTA2 (2.1e-51), Dynamin_N (2.3e-31), and Dynamin_M (2.7e-20) domains,
- a 271-residue protein with a FTA2 domain (1.2e-20),
- a 405-residue protein with FTA2 (1.6e-42) and HlyIII (3.7e-08) domains,
- a 864-residue protein with 4 PDZ domains (6.3e-06 to 3.4e-17),
- a 254-residue protein with a Homeodomain domain (3.4e-14),
- a 569-residue protein with Hsp70 (6.6e-42) and MreB/Mbl (2.4e-10) domains,
- a 138-residue protein with an NUDIX domain (1.2e-14),
- a 181-residue protein with an Scm3 domain (2.2e-14),
- a 184-residue protein with a CinA domain (1.6e-46),
- a 268-residue protein with a Helo domain (1.7e-17),
- a 276-residue protein with an NUDIX domain (2.3e-05).

Note that the two proteins with Hsp70 MreB/Mbl domains and the one with a CinA domain had few homologs in other fungi, where CinA is thought to be involved in the process of transformation.

